# Interpretable prioritization of splice variants in diagnostic next-generation sequencing

**DOI:** 10.1101/2021.01.28.428499

**Authors:** Daniel Danis, Julius O.B. Jacobsen, Leigh Carmody, Michael Gargano, Julie A McMurry, Ayushi Hegde, Melissa A Haendel, Giorgio Valentini, Damian Smedley, Peter N Robinson

## Abstract

A critical challenge in genetic diagnostics is the computational assessment of candidate splice variants, specifically the interpretation of nucleotide changes located outside of the highly conserved dinucleotide sequences at the 5′ and 3′ ends of introns. To address this gap, we developed the Super Quick Information-content Random-forest Learning of Splice variants (SQUIRLS) algorithm. SQUIRLS generates a small set of interpretable features for machine learning by calculating the information-content (IC) of wildtype and variant sequences of canonical and cryptic splice sites, assessing changes in candidate splicing regulatory sequences, and incorporating characteristics of the sequence such as exon length, disruptions of the AG exclusion zone, and conservation. We curated a comprehensive collection of disease-associated splice-altering variants at positions outside of the highly conserved AG/GT dinucleotides at the termini of introns. SQUIRLS trains two random-forest classifiers for the donor and for the acceptor and combines their outputs by logistic regression to yield a final score. We show that SQUIRLS transcends previous state of the art accuracy in classifying splice variants as assessed by rank analysis in simulated exomes and is significantly faster than competing methods. SQUIRLS provides tabular output files for incorporation into diagnostic pipelines for exome and genome analysis, as well as visualizations that contextualize predicted effects of variants on splicing to make it easier to interpret splice variants in diagnostic settings

## INTRODUCTION

Whole exome sequencing (WES) and whole-genome sequencing (WGS) are effective tools to diagnose Mendelian disorders. However, although the diagnostic yield of WES/WGS has improved from between 16-25% in early studies^1–3^ to around 35-60% currently,^4,5^ a substantial proportion of diagnostic cases remain unsolved. One reason is that the filtering and prioritization typically used by diagnostic WES/WGS software is not able to correctly classify some kinds of disease-causing variants. It can be difficult to correctly classify splice-altering variants, especially those deep within exons or introns.^6^ Variants that affect pre-mRNA splicing are documented to account for at least 15% of disease-causing variants.^7^ However, the true number may be substantially higher because of a historical ascertainment bias reflecting a selective focus on coding sequences in the pre-next generation sequencing (NGS) era and a continued interpretation bottleneck due to the difficulty of predicting the effects of variants on splicing. For instance, in the NF1 and ATM genes, studies have shown that ~50% of all disease-causing variants result in defective splicing.^8,9^ Recent results have shown that RNA-seq may be able to identify the diagnosis in up to ~30% of exome-negative cases,^10–13^ and a massively parallel assay suggested that up to 10% of all exonic variants, including missense and nonsense variants, may alter splicing.^14^ However, RNA samples may not always be available in the diagnostic setting, and the relevant genes and transcripts may not be expressed in tissues commonly assayed for RNA analysis such as blood and muscle. A typical diagnostic exome or genome can contain over 500 candidate splice-altering variants of unknown significance.^15^ Therefore, there is a pressing need for algorithmic approaches that can effectively prioritize splice variants in diagnostic next-generation sequencing. Additionally, the interpretability of predictions is important for integration of results into medical workflows.^16^

For brevity, we use the term ‘splice-altering variant’ (SAV) to refer to disease-associated DNA variants that result in splice alterations. SAVs can lead to a number of molecular defects including exon skipping, cryptic splicing, intron inclusion, leaky splicing, or the introduction of pseudo-exons into the processed mRNA.^17^ There are no general rules that allow one to interpret the effect of a variant based solely on the affected sequence context, but it is generally accepted that alterations of the canonical ±1 or ±2 splice sites are most likely to be pathogenic. This is reflected in the fact that the American College of Medical Genetics (ACMG) guidelines state that the location of a variant in these positions can be taken as very strong evidence of pathogenicity in genes where loss of function is a known mechanism.^18^ However, the natural donor and acceptor splice sites span much longer intervals that overlap the exon-intron boundaries. In addition, the branch point and polypyrimidine tract motifs as well as intronic and exonic splicing enhancers and silencers further modulate the strength of any given splice site. Variants in any of these sequences can reduce or abolish the ability of the spliceosome to recognize the splice site, leading to exon skipping or usage of cryptic splice sites. The sequence between the branch point and the 3’ splice site is generally devoid of AG dinucleotides and is called the AG□exclusion zone; variants that introduce an AG in this zone tend to be pathogenic.^19^ Additionally, variants in introns or exons can activate cryptic splice sites to the extent that they are preferentially utilized compared to wildtype splice sites. We will use the term ‘canonical’ SAV to refer to variants at the ±1 or ±2 splice sites, and ‘non-canonical’ SAV to refer to any other SAV.

While canonical SAVs are trivial to identify computationally, non-canonical SAVs are substantially more difficult to interpret. Numerous bioinformatics tools such as PolyPhen^20^ have been developed to assess pathogenicity of missense variants, but far fewer have been developed for non-canonical SAVs. Suggestive evidence exists that non-canonical SAVs might be a more common cause of Mendelian disease than is commonly appreciated.^9,19,21^. Several previous approaches to prioritizing SAVs are based on information theory analysis, which compares wildtype and alternate sequences to a matrix of negative logarithms of the frequencies of nucleotides in the positions of wild type splice sites.^22^ Maximum entropy modeling of splicing signals (MaxEnt) is a similar approach that additionally may include dependencies between nonadjacent as well as adjacent positions.^23^

Numerous algorithms have been presented for the prioritization of SAVs.^24–29^ Recently, machine learning methods surpassed previous state-of-the-art results in the prediction of pathogenic SAVs including sequence□based deep neural networks^30,31^ and gradient boosting trees.^15^ However, it is not straightforward to interpret the results of these methods. For instance, SpliceAI is a deep residual neural network that predicts whether each position in a pre-mRNA is a splice donor, splice acceptor, or neither; differences in the scores of wildtype and variant sequences can be used to predict pathogenicity of variants, but no information is provided by the algorithm as to what sequence features led to the prediction.^31^ This makes it challenging to use in a clinical setting, where explainability is essential for clinical decision making. S-CAP uses a gradient-boosting tree (GBT) classifier, with 29 features including predictions from a number of other algorithms; the results of the algorithm are presented as a single score that does not allow further interpretation.^15^

Here we present a new algorithm, Super Quick Information-content Random-forest Learning of Splice variants (SQUIRLS). SQUIRLS first scores variants according to associated changes in individual information content (ΔIC), changes in splicing regulatory elements (SREs), and several other features, followed by random forest classification. SQUIRLS was trained on a comprehensive dataset of 1,623 non-canonical SAVs. SQUIRLS prioritized more correct variants in the top five ranks, with substantially higher speed and interpretability than the previously proposed best performing methods.^15,31^ The results can be output with visualizations and assessments of each feature, allowing users to quickly identify the major abnormalities that led to the prioritization. SQUIRLS is an interpretable and fast machine-learning algorithm that assesses variants for potential effects on splicing. SQUIRLS was designed to perform well on difficult to classify non-canonical splice variants located outside of the nearly perfectly conserved AG/GT dinucleotides at the termini of introns. We believe that SQUIRLS will support improved and scalable diagnostic capability for clinical interpretation of splice variants identified by WES/WGS.

## METHODS

### Dataset of splice variants

We performed an extensive review of the scientific literature to curate a collection of 8,314 splice variants associated with Mendelian diseases. Candidates were derived from a review of ClinVar pathogenic mutations^32^ and a manual review of the medical literature. We included case reports, mutation updates, and review articles describing variants where splicing deleteriousness was supported by experimental evidence, such as minigene assay, site directed mutagenesis, or patient-derived RNA sample analysis. We also included cases where the proband’s phenotype corresponded to the phenotype of the Mendelian disease associated with the affected gene. Our review of ClinVar database focused on synonymous pathogenic mutations as well as on non-canonical SAV that overlap with canonical splice site regions. The variants are listed in the Supplemental Table 1. The curated variants were located on chromosomes 1-22 and chromosome X (minimum count per chromosome: 77 for chr21; maximum: 1339 for chrX) and were derived from a total of 4522 articles with PubMed ids. 4753 were assigned to the donor site, 3388 to the acceptor site, and 173 were not assigned to a specific site. Variants from 1080 genes were included, with 370 genes with just one SAV, 401 genes with 2-5 SAVs each, 233 genes with 6-20 SAVs, 50 genes with 21-50 SAVs, and 26 genes with over 50 SAVs.

### Dataset of non-deleterious variants

We prepared a collection of 73,203 presumed non-deleterious variants from the ClinVar database.^32^ After downloading the VCF file released on Nov 11, 2019 from the ClinVar FTP site, we selected variants where both the *wt* and *alt* alleles were shorter than 50bp, the clinical significance of the variant was classified as either *benign* or *likely benign*, and the variant was located in coding region of a gene or distance from the closest exon was less than 100bp. Each non-deleterious variant was assigned to a donor and/or acceptor site, depending on distance to the site.

### Engineering of the splicing features

We developed a set of numeric features to discriminate splicing pathogenic variants from the neutral variants. The features can be separated into three groups: a) information content features, b) features representing the sequence context, and c) variant site features.

The first group of features is related to the *individual information content* of the affected sequences.^22^ We compute the individual information content of the closest canonical splice sites and the maximum *information content* of the surrounding *wt* sequence to model the inherent potential of the *wt* sequence for abnormal splicing. Then, *the differential information content-based* features represent an estimate of changes to free energy of binding of spliceosome components of pre-mRNA induced by the *alt* allele.

The sequence context features include length of the closest exon and the offset (distance in nucleotides) to the closest canonical splice sites to capture potential positional dependencies. The two remaining features of this group identify variants that introduce an AG dinucleotide into the AG exclusion zone (the sequence between the branch point and the 3’ splice site that is devoid of AGs). In our implementation, the AGEZ is defined to be positions −50 to −3, although biologically, the branchpoint is located between −18 and −40 (and not reliably identifiable computationally).

The variant site features are calculated for the nucleotides that are altered by the variant. We use ESRSeq^33^ and SMS^34^ to assess changes to splicing regulatory element sequences that are associated with exon skipping and inclusion and may be related to functional elements such as exonic splicing enhancers for which currently no sensitive and specific sequence motifs are available. *phyloP* evolutionary conservation scoring^35^ reflects whether the nucleotide or nucleotides altered by the variant are under natural selection against a background of neutral evolution.

In the next section we describe more in detail the construction of the features based on the information content of the sequences. Table 1 provides an overview of features, and the following sections provide additional details.

**Table 1.** Features used to discriminate deleterious splice variants from splicing neutral in SQUIRLS. We used 7 features to train site specific random forest classifiers for donor variants, and 9 features to train the classifier for acceptor variants. Note that phyloP is used by both splice donor and acceptor classifiers. *R_i_* - information content of a nucleotide sequence in bits, *AGEZ* (AG-exclusion zone) - canonical acceptor site region located between the authentic 3’ss AG and the branch point that is generally devoid of AG dinucleotides.

|  | **Splicing feature name** | **Description** |
| --- | --- | --- |
| Donor | Donor offset | Distance to the exon/intron border of the closest donor site. The number is negative if the variant is located upstream from the border. |
| | $R_i$ can ref | Information content ($R_i$) of the closest canonical donor site. |
| | max $R_i$ cryptic donor window | Maximum $R_i$ of sliding window of all 9 bp sequences that contain the *alt* allele. |
| | $\Delta IC_{can}$ | Difference between $R_i$ of *ref* and *alt* alleles of the closest donor site (0 if the variant does not affect the site). |
| | $\Delta IC_{crypt}$ | Difference between max $R_i$ of sliding window of all 9 bp long sequences that contain the *alt* allele and $R_i$ of *alt* allele of the closest donor site. |
| | $\Delta IC_{next}$ | Difference between $R_i$ of the closest donor and the downstream (3') donor site (0 if this is the donor site of the last intron). |
|  | phyloP | Mean phyloP score of the reference nucleotides altered by the variant, where phyloP denotes conservation scoring calculated by PHAST package for multiple alignments of 99 vertebrate genomes to the human genome<sup>35</sup> |
| Acceptor | Acceptor offset | Distance to the exon/intron border of the closest acceptor site. The number is negative if the variant is located upstream from the border. |
| | $\Delta IC_{can}$ | Difference between $R_i$ of *ref* and *alt* alleles of the closest acceptor site (0 if the variant does not affect the site). |
| | $\Delta IC_{crypt}$ | Difference between max $R_i$ of sliding window applied to *alt* allele neighboring sequence and $R_i$ of *alt* allele of the closest acceptor site. |
|  | Exon length | Number of nucleotides spanned by the exon where the variant is located in (-1 for non-coding variants that do not affect the canonical donor/acceptor regions). |
|  | Creates 'AG' in AGEZ | 1 (True) if the variant creates a novel 'AG' dinucleotide in AGEZ, 0 (False) otherwise. |
|  | Creates 'YAG' in AGEZ | 1 (True) if the variant creates a novel 'YAG' trinucleotide in AGEZ where 'Y' stands for pyrimidine derivatives (cytosine or thymine), 0 (False) otherwise. |
|  | ESRSeq | Estimate of impact of random hexamer sequences on splicing efficiency when inserted into five distinct positions of two different minigene exons obtained by *in vitro* screening.<sup>33,36</sup> |
|  | SMS | Estimated splicing efficiency for 7-mer sequences obtained by saturating a model exon with single and double base substitutions (saturation mutagenesis derived splicing score).<sup>34</sup> |
|  | phyloP | See above. |

### Features based on the information content of the sequences

The core features used to train the splice donor and acceptor site models are based on information theory applied to the analysis of splice sites.^22^ First, to construct a matrix with frequencies of nucleotides occurring at different positions of the splice sites, we aligned wild-type sequences of exon/intron junctions of GENCODE basic gene annotation transcripts v32 (accessed at Oct 2019). We selected 49,821 protein coding transcripts with gene annotation source Havana and GENCODE confidence level ≦2, corresponding to transcripts supported by the highest amount of the experimental evidence.

Then, we grouped the transcripts by gene and identified genomic coordinates of unique exon/intron junctions, producing sets with 200,459 donor and 197,874 acceptor site coordinates. Next, we extracted ±80bp of the nucleotide sequence surrounding the sites and we subsequently aligned the sequences by exon/intron junction coordinate. After alignment, we calculated a matrix, *F^4×m^* where 4 refers to the number of different types of nucleotides and *m* to the length of the sequences. Each element *f*(*b, l*) of the matrix *F* represents a probability of observing base *b* ∈ {*A, C, G, T*} at position *`l`* within the aligned sequences (Figure S4). Finally, we created an information weight matrix *`R`_iw_* grounded in the concept of “decrease in surprisal”^37^ to model a splice junction by the equation

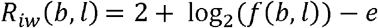

where *e* is a sample size correction factor for the *`n`* sequences at position *`l`*.^38^ The *R_iw_* matrix represents the sequence conservation of each nucleotide within the binding site, measured in bits of information. After checking for background noise, we determined the lengths of the donor and acceptor sites to be l_don_ = 9bp and l_acc_= 27bp (see Figure S4 for more details).

The *R_iw_* matrix can be used to calculate the individual information content *R_i_* of any nucleotide sequence *j* with length *m* as:

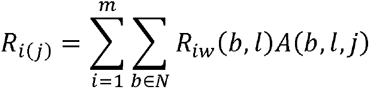

where *N* ∈ {*A,C,G,T*} is the set of nucleotides, and *A* is a *4* x *m* binary matrix that represents a one-hot encoding of the sequence *j:* the *A* matrix has only one 1 for each column while the rest of its column elements is set to 0. In effect, each base of the sequence “picks out” a specific entry of the matrix *R_iw_* and these entries are finally added to compute the information content of the sequence. In our setting, *R_iw_* is a weight matrix representing the splice junction, and the mean values of the *R_i_* distribution for the donor and acceptor sites, that represent the mean information of the sequences used to construct *R_iw_*, were 7.87 (donor) and 9.50 (acceptor) bits. The resulting *R_i_(j)* is related to thermodynamic entropy and the free energy of binding and can be used to compare sites with one another.^38^

### Training and test variant sets

We pooled the splice and neutral variants and then we annotated each variant with splicing features (Table 1) and additional metadata, including label (deleterious or neutral), gene symbol, transcript accession ID, and cytoband. Next, we split the variants into train and test sets by applying a “cytoband-aware” hold-out scheme: we randomly chose 10% (67) of the total number of 676 cytobands, and we put the variants contained in these cytobands into the test set. The variants located in the remaining 90% (609) cytobands were used for training (Fig. S5). The cytoband-based scheme was designed to minimize bias resulting from distinct variants located in the same gene being used for both training and testing. Then, we partitioned the training variants into two subsets consisting of either donor or acceptor-affecting variants, based on curation metadata or vicinity to one or the other splice site. We removed 6008 canonical SAV variants from the training set, since we aimed to optimize the classifier for non-canonical SAVs. We tested SQUIRLS using both subset of non-canonical SAVs as well as the entire set.

### Training of the SQUIRLS model

SQUIRLS is a “paired ensemble” model that predicts the potential of a variant to alter the splicing pattern of an overlapping transcript. The model consists of two random forest classifiers^39^ trained individually on either the donor or the acceptor variant subset. If features are missing for a data point, they are replaced by the median value prior to random forest analysis.

To train the classifiers and perform model selection, we ran 50 iterations of randomized search cross-validation. In each iteration we randomly sampled hyperparameter values from pre-defined parameter distributions and performed 10-fold cross-validation on the training set. Each cross-validation step included calculation of the following performance metrics: balanced accuracy, precision, recall, and F1 scores.

We selected the hyperparameters that produced the model with the highest sensitivity (recall) and we subsequently retrained the donor and acceptor classifiers on the whole variant subset.

Most of the machine learning methods used to identify potential pathogenic variants report predicted deleteriousness/pathogenicity estimates as a number in the range [0,1], where higher scoring variants are more likely to be deleterious.^40,41 42^ In addition, thresholds for assigning variants into discrete classes (e.g. neutral and deleterious) while obtaining the desired specificity or sensitivity are available for most of the methods. In a random forest, probability estimates for a class can be calculated as the proportion of the forest’s decision trees that voted for the class. To find the class probability threshold that attains the best separation of splice and neutral variants, we used the value that maximized the informedness criterion (Youden’s J statistic).

To learn the final SQUIRLS score, we applied a meta-learning approach by stacking a learning machine on top of the two random forests.^43^ More precisely, we trained a logistic regression model (LR) from the raw scores computed by the two random forests, to automatically learn how to better combine their output.

For model training and evaluation, we used random forest, logistic regression, and imputer implementations provided within the Scikit-learn framework.^44^ For the SQUIRLS application and library, we wrote a custom implementation of the imputer, random forest, and logistic regression. The implementation is available in the SQUIRLS source code repository (Web resources).

### Model testing, validation, and comparison with other splicing pathogenicity algorithms

To obtain the unbiased performance estimate for SQUIRLS scores, we computed pathogenicity estimates for the test set variants and then we performed ROC and precision-recall analysis. We used the thresholds and evaluated classification accuracy.

We compared the SQUIRLS scores with other algorithms that are used for prioritization of splice variants. We chose two algorithms designed to assess splice variants that performed well in recently published analyses (SpliceAI^31^ and S-CAP^15^), an older well-established method (MaxEntScan^23^), and an algorithm that is commonly used for variant prioritization in WES/WGS experiments even though it was not specifically designed for analysis of splice variants (CADD^45^). To evaluate the ability of all algorithms to discriminate between the neutral and the splice variants, we calculated predictions for variants and constructed ROC and PR curves.

### SpliceAI

`SpliceAI` provides four delta scores for each variant where the maximum score denotes a probability of the variant being splice-altering.^31^ In order to evaluate `SpliceAI` performance, we precalculated the delta scores for variants in our dataset. We used version 1.3.1 (accessed on April 25 2020 at *https://github.com/Illumina/SpliceAI*) with the -`M True` option to mask scores representing annotated acceptor/donor gain and unannotated acceptor/donor loss. We chose the maximum value to perform ROC and PR evaluation. We benchmarked SpliceAI runtime performance using the Python package `spliceai` v1.3.1 available at PyPi. The runtime of `spliceai` for a single VCF file with ~100,000 variants is roughly one day, so we benchmarked `spliceai` on VCF files subsampled to 5,000 variants only.

### S-CAP

The `S-CAP` algorithm provides splicing-specific pathogenicity scores calculated using gradient-boosting tree (GBT) algorithm.^15^ The algorithm consists of six GBT predictors, one predictor for each of six author-defined regions relative to the splice site. The authors provide a VCF file with precomputed scores for all possible single nucleotide variants in the splicing region. There are two score types: *raw* score is the output of the corresponding GBT, and *sensitivity* score which is a transformed *raw* score to make it directly comparable with scores of the other regional predictors. We used both *raw* and *sensitivity* scores for the ROC and PR evaluation.

### MaxEntScan

`MaxEntScan` is a framework that employs the maximum entropy principle for building a model *m* that represents a particular sequence motif, including mRNA splice sites.^23^ During the building phase, a collection of aligned sequences is used to estimate the maximum entropy distribution and a set of constraints. Using this approach, the authors built and evaluated multiple maximum entropy models. For our comparison, we chose the models that yielded the highest AUCs (*m_me2×5_* for the donor and *m_me2×3 for the_* acceptor site), as described in the `MaxEntScan` manuscript.

In order to allow `MaxEntScan` to be compared with SQUIRLS, we created a set of rules for constructing nucleotide snippets *j_wt_* and *j_alt_* to be scored by the appropriate `MaxEntScan` model *m*. For each variant, we considered four situations:

(i)the variant disrupts the canonical donor site, (ii) the variant activates a cryptic donor site, (iii) the variant disrupts the canonical acceptor site, and (iv) the variant activates a cryptic acceptor site.

For situations (i) and (iii), we prepared sequence snippets *j_wt_* and *j_alt_* for the canonical sites and we calculated the final score *Δ_MES_* as *Δ_MES_* = *m(j_wt_) - m(j_alt_)*. For situations (ii) and (iv), we calculated a score vector **s** for the sliding window of all *n*-bp sequences *j_wt_* or *j_alt_* that contain the *wt or alt* alleles. Then, the final score was computed as *Δ_MES_* = *max*(**s**_alt_) - *max*(**s**_wt_). After calculating *Δ_MES_* for all four situations, we used the maximum value as the final pathogenicity estimate for ROC and PR analysis.

### Combined Annotation Dependent Depletion

Combined Annotation Dependent Depletion (`CADD`) estimates the deleteriousness of variants by integrating multiple annotations into a single score.^45^ The score is applicable across diverse variant functional categories, including variants affecting mRNA splicing. For comparing `CADD` with SQUIRLS, we downloaded TSV files with PHRED-scaled pathogenicity scores precalculated for all possible SNVs and INDELs built by the model v1.4 (accessed on November 20, 2019). For each variant, we transformed the PHRED score *x* into [0,1] by applying 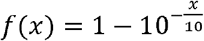. If the score was not available, we considered the variant to be benign (pathogenicity=0.0). The transformed scores were used for ROC and PR analysis.

### Implementation

We designed multiple optimizations to achieve fast runtime performance. SQUIRLS fetches all data required to evaluate a variant’s effect on the overlapping transcripts in a single I/O lookup and all the subsequent operations are performed in memory. An additional performance increase is achieved by limiting the number of splicing features and by exploiting inherent parallelism of the random forest, which can be distributed across multiple CPU cores. The source code of SQUIRLS and a standalone “executable JAR” file are available for download from the GitHub repository (Web resources).

## RESULTS

SQUIRLS is designed to predict variants associated with splice defects from exome or genome sequencing data. All variants that overlap transcripts are evaluated for potential effects on splicing including both variants at the canonical donor and acceptor sequences as well as other exonic and intronic variants that could generate cryptic splice sites or otherwise alter normal splicing (Fig. 1A). SQUIRLS evaluates the effect of variants with respect to all transcripts that overlap the variant. The output visualizations and tabular assessments are designed for human consumption and can also be used to output a VCF file with annotations of the predictions of relevant splice variants for use in larger bioinformatic pipelines for diagnostic genomics.

**Figure 1.).**
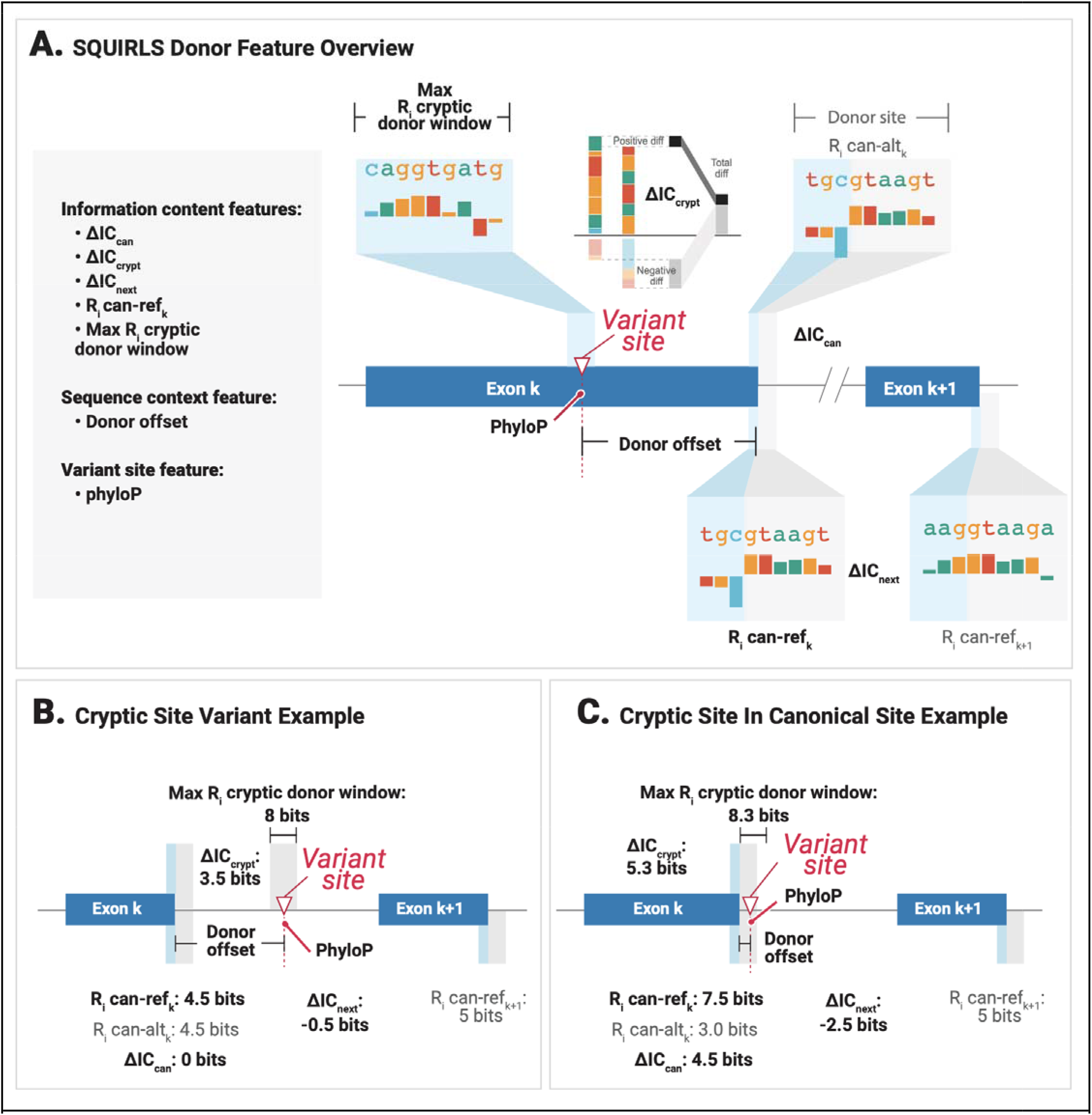
mRNA splicing and sequence logos/walkers. **A)** The figure shows an intron and the corresponding canonical splice donor and acceptor sites, which are represented as logos, where the letters representing the sequence are stacked on top of each other for each position in the splice site. The height of each letter is proportional to its logarithmic frequency at the position, and the height of the entire stack is adjusted to indicate the information content of the sequences at that position. **B)** Individual sequence information (*R_i_*) for a wildtype splice donor sequence of *CHRNE* and for the corresponding sequence with the variant NM_000080.3:c.917G>T:p.(R306M). c.917G>T is located at the last (3’ most) position of an exon and although it is predicted to lead to a missense change, it reduces the strength of the donor sequence and leads to skipping of the affected exon.^46^ The sequence walker representations as introduced by Rogan and colleagues^22^ are shown for the wildtype and variant sequences. Sequence walkers display nucleotides that represent favorable contacts to the spliceosome and a test sequence by letters that extend upwards and positions that are predicted to make unfavorable contacts are shown by inverted letters. **C)** SQUIRLS introduces a new graphical representation in which a bar chart is used to show the degree to which a sequence “matches” the donor or acceptor model. The height of the bars is calculated in the same way as for the height of the letters in the sequence walker. Positions that are changed by a variant are displayed such that the original nucleotide is shown as an outline (the “g” in this example) and the variant (alternate) base is shown filled. **D)** The variant reduces the *R_i_* from 7.6 to 4.0 bits. Changes in *R_i_* are referred to as ΔIC. SQUIRLS calculates ΔIC in several contexts (Fig. 2).

### Overview of the algorithm

SQUIRLS first calculates a set of numerical features for each variant/transcript pair. The features include changes in information content between reference and alternate alleles (**Fig. 1**), changes in SREs, distances from the canonical splice sites, and a measure of evolutionary conservation. The features were chosen to be interpretable by humans (**Table 1, Fig 2–3**). The features are used as input for a pair of random forest classifiers specialized in computing site-specific splice scores for donor and acceptor sites. The algorithm then uses logistic regression to transform the scores into the final SQUIRLS score that estimates the probability of the variant in question being a splice variant.

**Figure 2.**
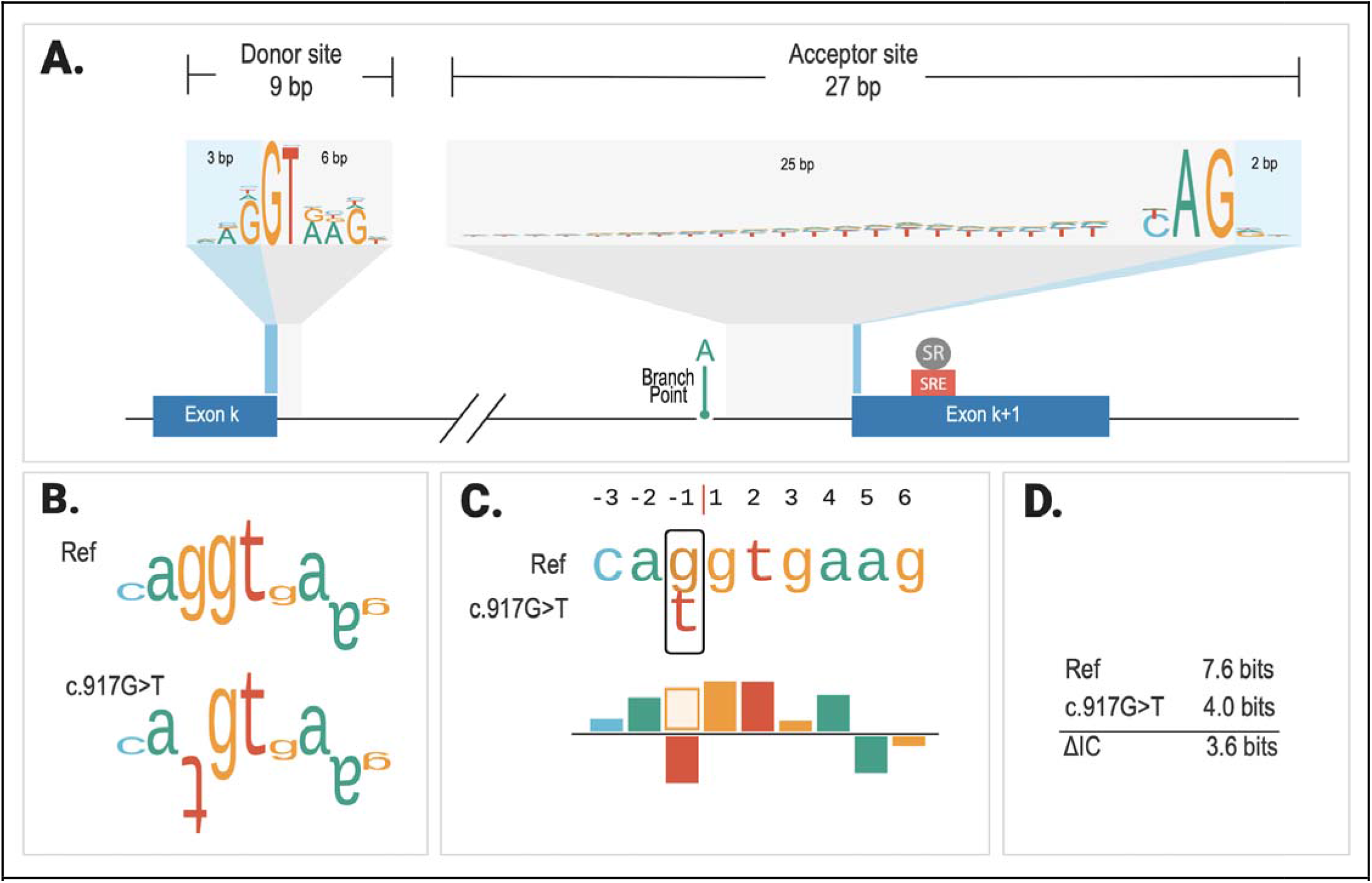
Calculate of changes in the individual information content (**A) Donor site.** SQUIRLS calculates ΔIC_can_ as the difference in the *R_i_* in between the reference and alternate sequence of the canonical donor sequence (Fig 1A). If the variant is located outside of this sequence, ΔIC_can_ =0. SQUIRLS evaluates the potential of variants to create cryptic splice sites using a sliding window approach (Methods). ΔIC_crypt_ is calculated by subtracting the *R_i_* of the reference sequence from that of the alternate sequence. Finally, the difference between the *R_i_* of the wildtype donor site is compared with that of the donor site of the following exon (ΔIC_next_), because differences in splice site strength can be predictive of exon skipping.^14^ **B) Acceptor** ΔIC_can_ and ΔIC_crypt_ are calculated in an analogous fashion. The random forest for acceptor variants does not use ΔIC_next_ as our initial analysis showed that it did not boost classification performance. See Table 1 for information about other features.

**Figure 3.**
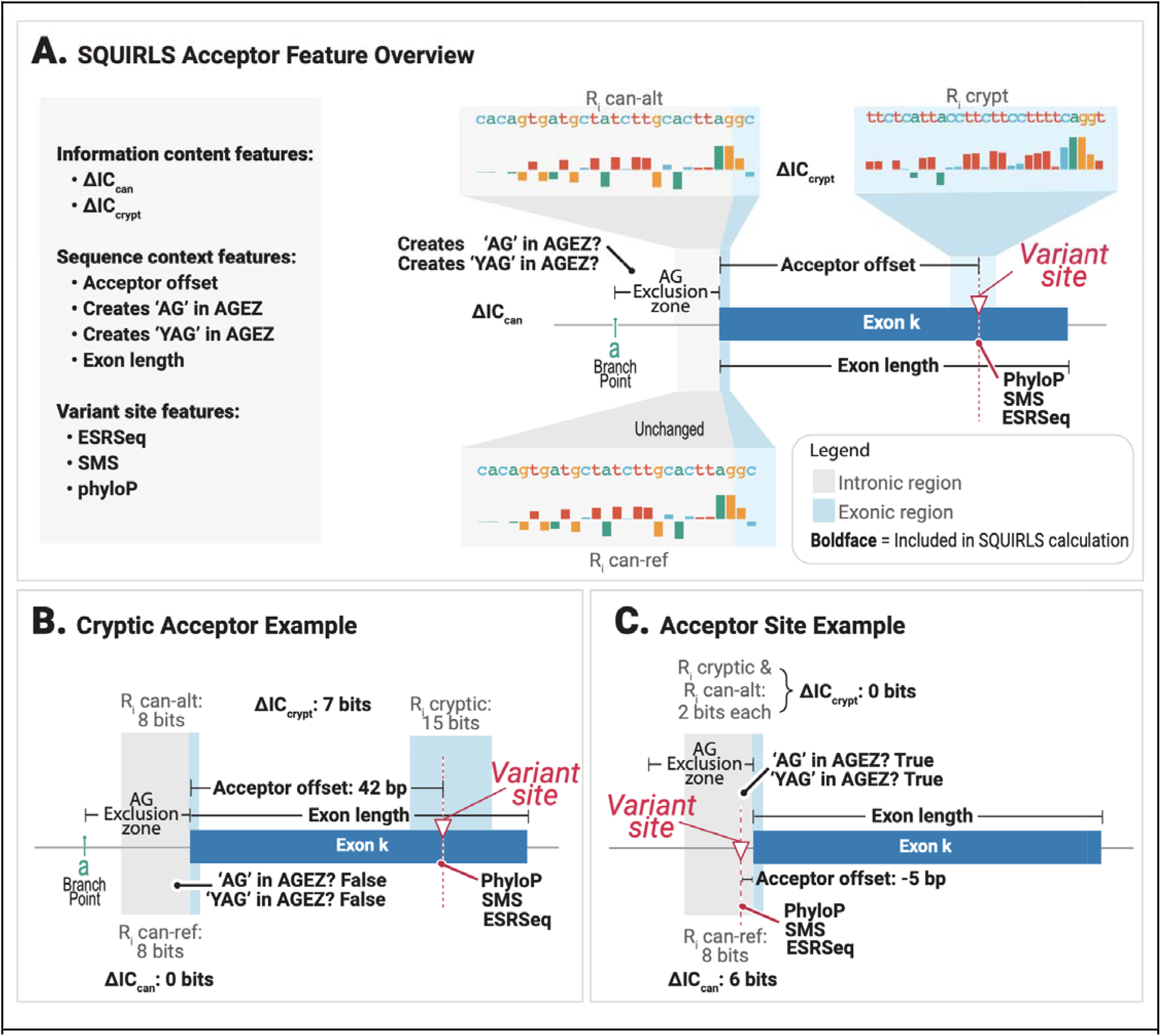
Features are extracted and generated for the donor and acceptor sites and leveraged for random forest learning, whose predictions are calibrated by logistic regression to provide the final SQUIRLS predictions. **A)** SQUIRLS calculates 7 features to evaluate variant impact on the donor site. The individual information content (R_i_) of the reference and alternate canonical splice site and of the donor site in the following exon (Exon *k+1*) are calculated and used to determine the difference in information content between the reference and alternate canonical splice site (ΔIC_can_), the difference between the best candidate cryptic splice site and the alternate sequence of the canonical splice site (ΔIC_crypt_) and the difference between the donor site at exon *k* and *k+1* (ΔIC_next_). See Table 1 for information about other features. **B)** In this example, a variant in intron k creates a cryptic splice site with 8 bits, which is greater than in the individual information of the canonical splice site (4.5 bits), so ΔIC_crypt_ =3.5 bits. The variant does not change the sequence of the canonical splice site, so ΔIC_can_=0. The the individual information of the donor site of the next exon has 0.5 more bits greater than that of exon k, so ΔIC_next_ =−0.5 bits. **C)** In this example, a variant in the canonical splice site (e.g., the +5 position) reduces the strength of the canonical splice site from 7.5 to 3.0 bits and simultaneously creates a novel cryptic site with an individual information content of 8.3 bits. An example of this is the variant NM_000314.7(PTEN):c.253+2T>C, which alters the canonical splice site and simultaneously changes the sequence of a cryptic splice site located 3 nucleotides downstream, resulting in the inclusion of 4 intronic nucleotides in the variant mRNA.^47^

### A dataset of non-canonical splice variants

We performed a comprehensive review of scientific literature to curate a dataset of splice variants associated with Mendelian diseases. In total, we collected 8,314 splice variants as well as 73,203 variants classified as benign or likely-benign variants from ClinVar (Table 2, Supplemental Table 1).^32^ The distribution of the variants with respect to the donor and acceptor splice site is shown in **Fig. 4**.

**Figure 4.**
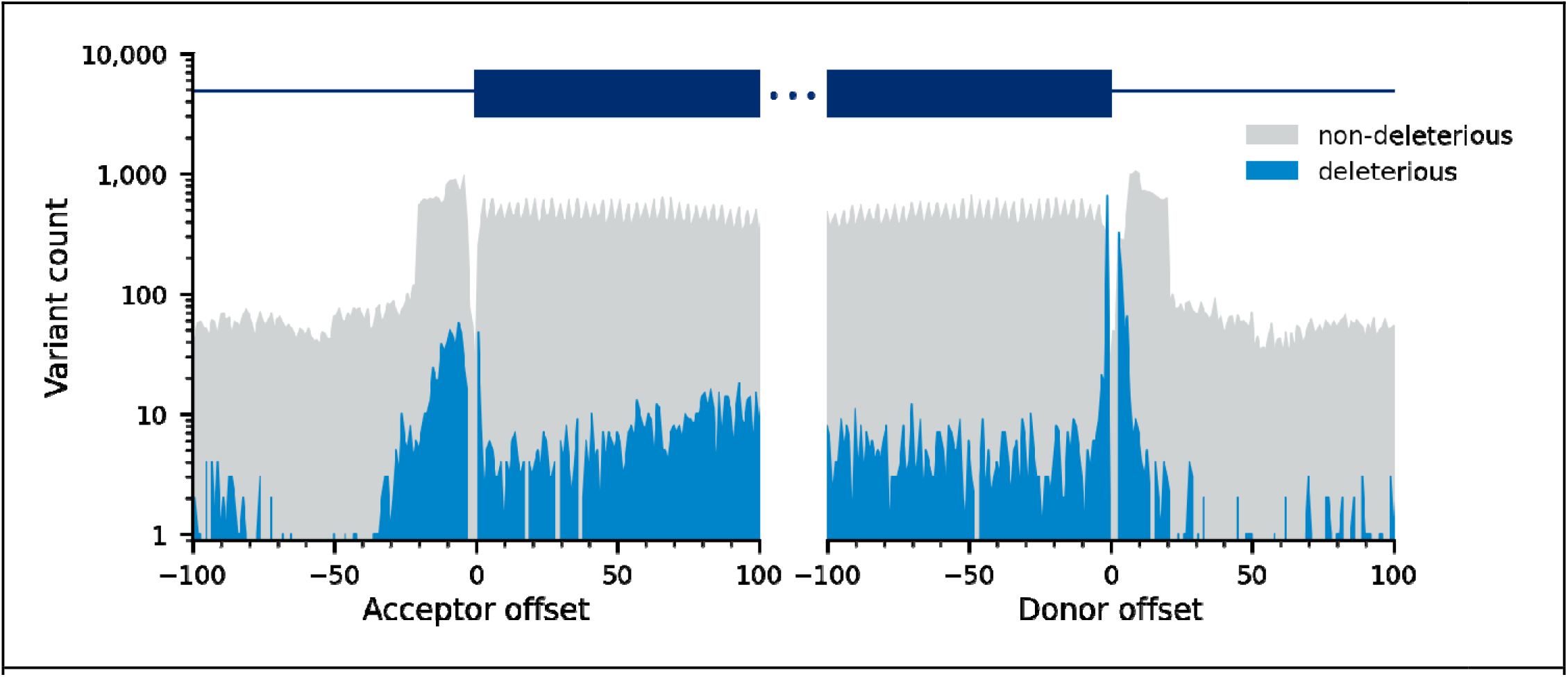
Distribution of non-canonical deleterious SAVs and non-deleterious variants used for training SQUIRLS. The figure shows the distribution of variants used for training SQUIRLS on a logarithmic scale. The position with respect to the nearest acceptor or donor intron/exon boundary is shown.

**Table 2.** Summary of the variant dataset. We created a collection of splice variants by curating literature. During curation, we recorded metadata regarding the variant pathomechanism and the observed outcome. Based on the outcome, we categorized the variants into two major groups: a) variants disrupting canonical splice sites and leading to activation of a cryptic splice site, or to exon skipping, and b) variants that activate cryptic splice site outside of the canonical splice site. 73,203 neutral variants were used as negative training examples.

| Outcome | Donor | Acceptor | Total |
| --- | --- | --- | --- |
| Cryptic site creation | 150 | 204 | 354 |
| Canonical site disrupted | 4,701 | 3,242 | 7,943 |
| Other | 7 | 10 | 17 |
| **Total** | **4,858** | **3,456** | **8,314** |

In order to prepare the variant dataset for training of machine learning models, we split the dataset into training and test sets. We used a “cytogenetic band-aware” method that ensures that variants affecting the same gene are either used for training or testing, but not both, since nearby variants may share similar features which might bias the results. This way we randomly partitioned the splice and non-deleterious variants into training (609 cytobands, ~90%) and test (67 cytobands, ~10%) sets, consisting of 70,617 and 10,901 variants (Fig. S5).

Then, we assigned the training set variants to either donor or acceptor sites, based on the curation metadata or distance to the closest splice site. The training set was further narrowed down by removing 6,008 canonical SAVs, yielding the final training set consisting of 1,623 deleterious noncanonical SAVs and 62,986 non-deleterious variants. We chose to train SQUIRLS on non-canonical SAVs but note that SQUIRLS also displays state of the art performance in the (relatively simple) classification task of predicting deleteriousness of canonical SAVs.

### Selection of interpretable features for machine learning

We trained two site-specific random forest classifiers to separate splice variants from neutral variants, one for the donor variants and the other for the acceptor variants. During training, we used random search hyperparameter optimization^48^ and 10-fold cross-validation to evaluate different combinations of 21 splicing features and learning parameters, to select the combination that provides classifiers with the highest area under receiver operating characteristic curve (AUROC) and precision-recall scores. The final set of 15 features included features based on information content, changes in candidate 6/7-mer SRE motifs, evolutionary conservation of the variant position, and distance from the closest splice sites (Fig. 2A-B, Supplemental Fig. S1, Table 1). After selecting the best-performing features and learning parameters, we trained the final site-specific classifiers using the entire training set.

The donor and acceptor scores are calculated for all variants. The ranges and thresholds of the acceptor and donor scores are, however, different (Figure S2A), which precludes direct integration of the site-specific estimators into variant prioritization frameworks. To combine the donor and acceptor estimators into a single measure, we introduced a meta-learning approach using logistic regression as the last step of our algorithm. We calculated site-specific deleteriousness estimations for all training variants and we subsequently used the site-specific estimates to obtain logistic regression parameters that provide the best predictions (splice deleterious=1, neutral=0). The final SQUIRLS score is the output of the logistic function, integrating the raw scores into a single measure with range [0,1].

### Performance evaluation and comparison with other methods

We evaluated SQUIRLS using a test set consisting of 808 splice variants (213 non-canonical SAVs) and 10,092 neutral variants (10,068 non-canonical SAVs) that were not used for training. After calculating SQUIRLS scores for all variants, we assessed diagnostic utility by creating receiver operating characteristic (ROC) and precision-recall (PR) curves, as well as calculating the area under the ROC (AUROC) and the average precision (AP).

SQUIRLS achieved an AUROC of 0.91 and an AP of 0.62 on a test set consisting only of non-canonical SAVs (Figure 5). Although SQUIRLS does not use canonical (±1,2) SAVs for training, it achieved an AUROC of 0.97 and an AP of 0.88 on a dataset that included both canonical SAVs and non-canonical SAVs (Figure S3). These results show that SQUIRLS can accurately identify both easy (canonical) and difficult to assess (non-canonical) SAVs.

**Figure 5.**
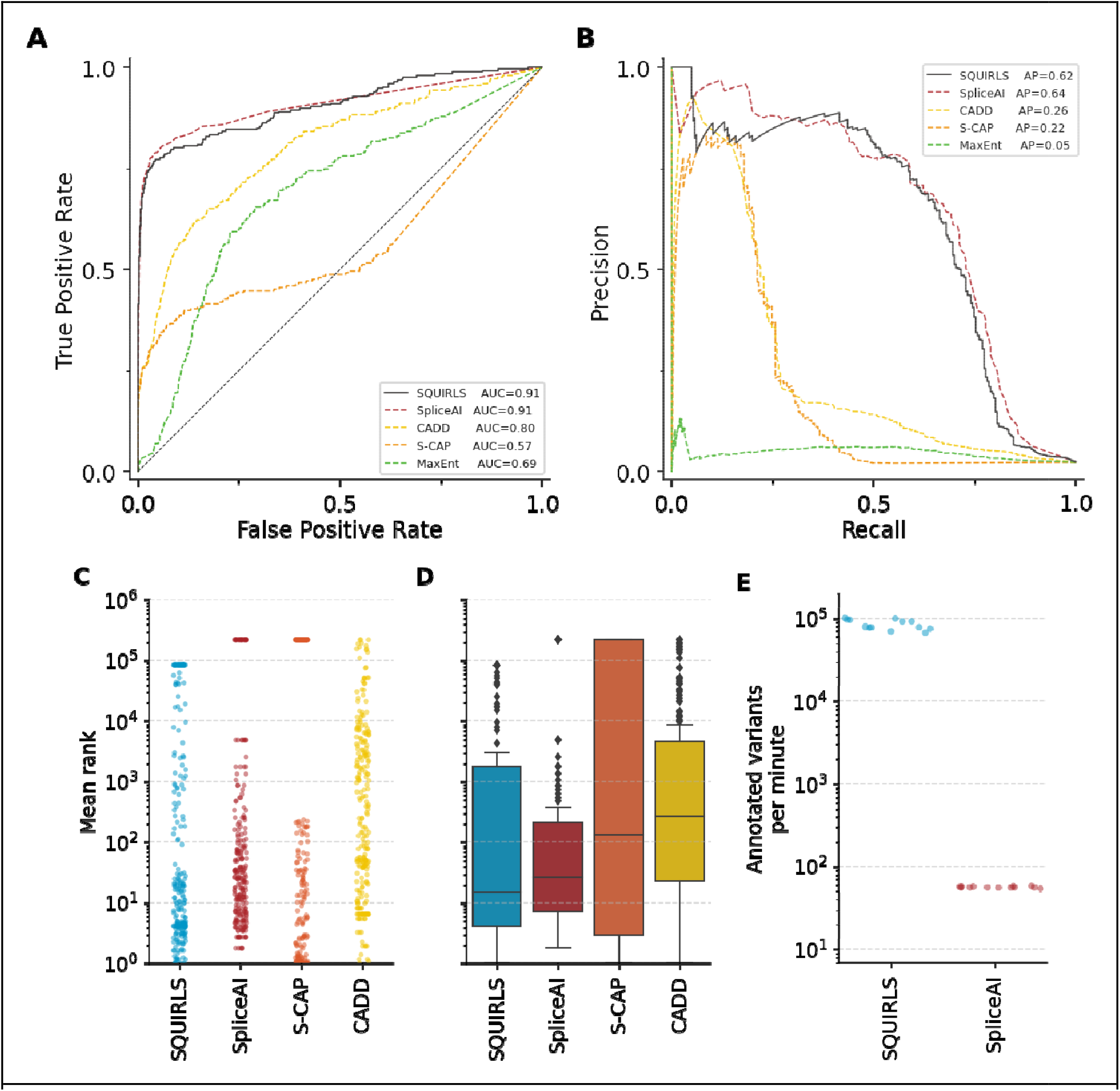
Performance of SQUIRLS, SpliceAI, S-CAP, CADD, and MaxEnt on non-canonical SVs. **A)** Receiver operating characteristic curves indicate that SQUIRLS and SpliceAI achieve comparable performance. **B)** Precision-recall curves show that SQUIRLS and SpliceAI are able to find the most of the true splice variants, while maintaining high precision. **C)** Mean ranks of splice variants among variants from 13 simulated exome sequencing runs. **D)** Mean ranks box plot. The horizontal line of each box indicates the median, box borders indicate positions of the 1^st^ and the 3^rd^ quartile, and the whiskers indicate 1.5x the interquartile range **E)** Comparison of algorithm runtimes for SQUIRLS and SpliceAI. We recorded the time required for analysis of 13 VCF files containing 87,000-107,000 variants. The figure shows the annotation speed that was achieved on a consumer laptop. We could not compare the performance of S-CAP and CADD, since they provide precomputed predictions as tabular files. Therefore, the annotation speed is only dependent on a package used to query the tabular file (e.g. tabix).

We then compared SQUIRLS to four state-of-the-art methods for assessing the pathogenicity of candidate splice variants: *SpliceAI,^31^* a deep residual neural network that predicts whether each position in a pre-mRNA transcript is a splice donor, acceptor, or neither, and *S-CAP,^15^* a gradient-boosting tree approach that provides splicing-specific pathogenicity scores. Moreover we compared SQUIRLS to *MaxEntScan,^23^* a well-established tool employing maximum entropy principle to model splicing motifs, and to *CADD,^45^* a framework that integrates diverse genome annotations into a single quantitative score to estimate deleterious effect of arbitrary variants and hence not specific for splice variants.

We obtained predictions for variants in the test dataset and constructed ROC curves and PR curves. SQUIRLS and SpliceAI achieved the best AUROC and AP on our test set, largely outperforming the other methods (Figure 4, Supplemental S3).

To further evaluate the expected performance of SQUIRLS in real-life scenarios, we developed a simulation strategy based on 13 VCF files generated by exome sequencing of individuals unaffected by a Mendelian disease. In the simulation, we added a single splice variant to each of the 13 VCF files, then we predicted pathogenicity for all variants, and subsequently ranked the variants according to predicted pathogenicity. Finally, we calculated the rank of the added splice variant averaged over the 13 VCF files.

In order for a prioritization method to be useful, it needs to place causal variants near the top of the list (“on the first page”) such that the causal variant is discoverable during the clinical interpretation. SQUIRLS achieved the best performance, placing 35% of splice variants within the top 5 positions, 50% of splice variants at rank 14 or below (median rank). The second-best method, SpliceAI, achieved a median rank of 25 and the third best method, S-CAP, achieved a median rank of 114 (Figure 5 C and D, S6).

### SQUIRLS enables rapid prioritization of arbitrary variants

With an ever increasing availability of sequencing data, computationally expensive algorithms may quickly become a bottleneck in the sequence data analysis. Precalculating pathogenicity scores for each genome position and storing the predictions in sorted and compressed tabular file or also using parallel hardware devices (e.g. graphics processing unit, GPU) are workarounds commonly used for computationally expensive algorithms. In contrast with SNVs, this approach does not work well for multi-nucleotide variants or INDELs, as the number of possible ref/alt allele combinations grows exponentially with increasing variant length. Then, storing pathogenicity prediction for each combination quickly becomes infeasible. Additionally, pre-calculated scores are not always available with respect to a particular transcript. To support pathogenicity prediction for an arbitrary genome variant at scale, the algorithm must be both efficient and easily portable to different computational platforms. SQUIRLS was designed to satisfy these requirements.

Apart from SpliceAI, SQUIRLS is the only tool in our comparison that directly annotates variants in a VCF file. S-CAP does not provide software that can analyze arbitrary variants, and a downloaded file with score mainly for single-nucleotide variants (SNVs) was used for the comparison. SQUIRLS annotates a VCF file containing 100,000 exome variants in roughly 1 minute on a consumer laptop, which is over 1000 times faster than SpliceAI (Fig. 5E). SpliceAI provides both a downloadable file with predictions for SNVs as well as an executable program that can analyze arbitrary variants. SQUIRLS was faster than all competitors except for the lookup of S-CAP predictions (Methods).

SQUIRLS is written in Java 11 and can be used both as a library, as well as a standalone command-line application (see online tutorial). The command line application is intended to be used with a Variant Call Format (VCF) file from exome or genome sequencing. The application generates output in multiple formats, including HTML report with figures and supporting information (see next Section), a tabular file with predictions, and an annotated VCF file that contains pathogenicity predictions with respect to all overlapping transcripts.

### SQUIRLS provides interpretable predictions

The majority of machine learning algorithms that are used as aids in variant prioritization work as black boxes. After making a prediction, the algorithms do not explain how the particular answer was made, which factors were considered, and the insights regarding the most likely molecular cause. When designing SQUIRLS, our motivation was to create an algorithm that is both accurate and interpretable. We addressed these goals by limiting features to a small set of biologically interpretable attributes (Table 1). SQUIRLS can output its results in three ways: (1) by adding annotations to the VCF file; (2) as a tab-separated values (TSV) file that can be easily incorporated into larger analysis pipelines; and (3) as an HTML file that presents the specific values calculated for each of the attributes relevant to a given variant in the context of visualizations that show the most important predicted effects. **Fig. 6** presents an example of the output produced by SQUIRLS for each candidate SAV.

**Figure 6.**
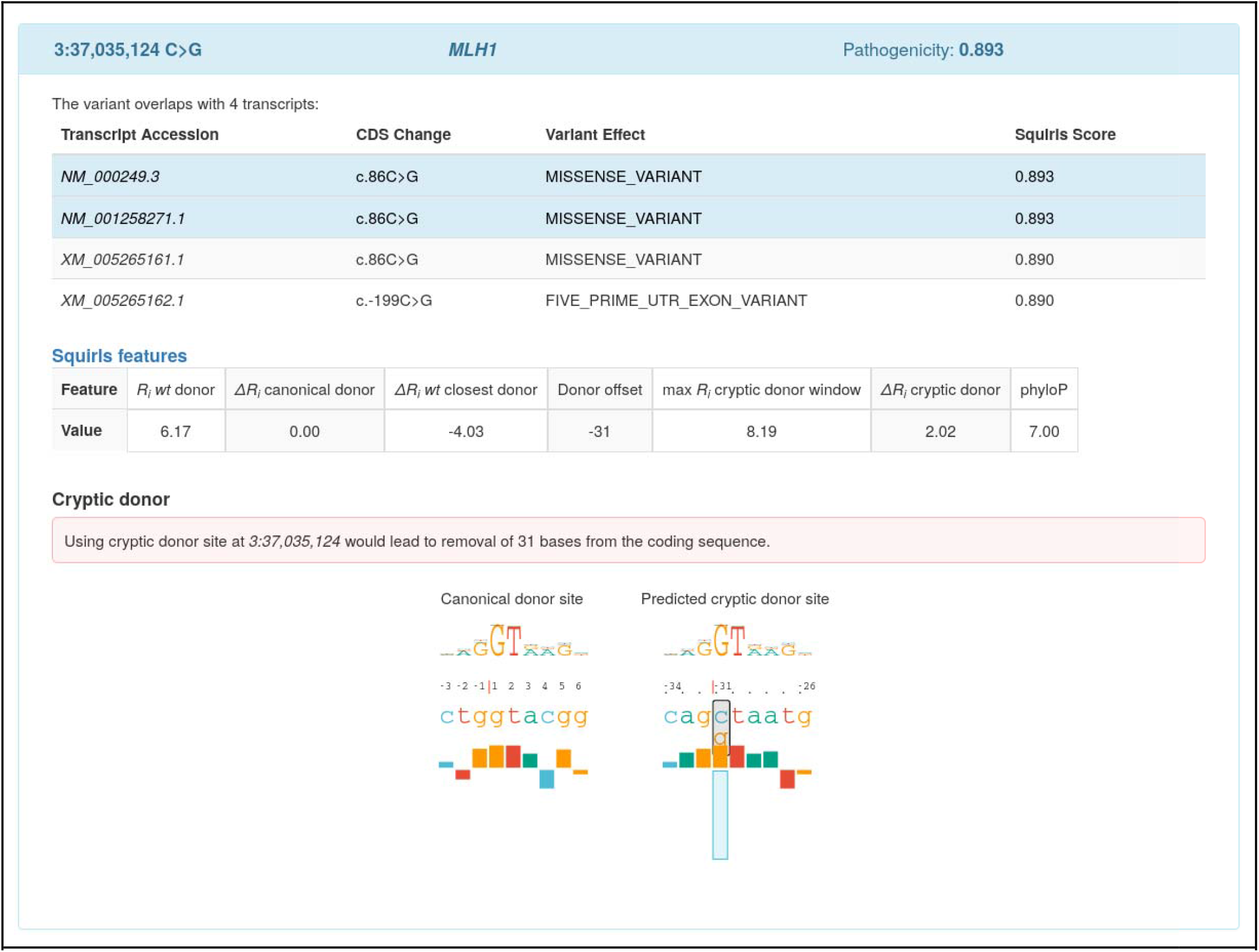
Screenshot of SQUIRLS HTML output. The variant NM_000249.3:c.86C>G generates a cryptic splice site in *MLH1.^49^* The variant is evaluated with respect to four overlapping transcripts and it is assigned maximum SQUIRLS score = 0.893. Transcripts with predicted maximum SQUIRLS score are highlighted in the table. The variant is located 31 bp upstream of the canonical site of exon 1 and it is predicted to create a cryptic donor site (*R_i_*=8.19 bits) which is stronger than the canonical donor (*R_i_*=6.17 bits) by 2.02 bits. Using the cryptic donor site would lead to removal of 31 bases from the coding sequence. Bar charts compare the canonical donor site with the predicted cryptic site. The bar chart shows that the variant replaces cytosine (blue rectangle) with a guanine (orange rectangle). The change is predicted to allow a more favorable contact between spliceosome and the *alt* allele, resulting in usage of the cryptic site and removing 31 bases from the coding sequence.

## DISCUSSION

In this work, we have presented SQUIRLS, an efficient and accurate algorithm for the prioritization of splice variants in exome or genome data. Our approach displays AUROC and AP performance that is comparable or better than that of previously published methods and is superior to these methods with respect to its ability to rank disease-associated variants within the long list of candidate splicing variants found in exomes. In contrast to previous methods, SQUIRLS was designed to leverage a small set of interpretable features and can provide visualizations of the predicted effects of variants on splicing that can help clinical interpretation.

To develop SQUIRLS, we focused on non-canonical splice variants. Canonical variants, defined as those that affect positions ±1 or ±2 of introns, are typically easy to interpret because variants at these positions only rarely do not deleteriously affect splicing. It has been substantially more difficult to develop algorithms that accurately classify splice variants at other positions. For this work, therefore, we performed extensive and detailed curation to identify non-canonical splice variants that are associated with Mendelian disease from the literature and from ClinVar. The resulting dataset, which to our knowledge is the largest of its kind, is freely available (Supplemental file 1). We developed a machine learning model using random forests and meta-learning techniques, whereby substantial preprocessing of sequence data is performed to generate a set of 15 features, using also information theory techniques to assess the information content of sequences that include splice variants. A meta learning approach based on stacked generalization is essential in this context to improve performance. Indeed a simple ensemble combination strategy based on averaging the raw scores computed by the random forests, or each random forest alone, worsens the overall performance (data not shown).

While SQUIRLS can be used on its own to specifically look for diagnostically relevant splice variants, it can also be easily used as a component of diagnostic exome/genome pipelines to improve recognition of causal splice variants. We optimized the classifier for high sensitivity to reduce the number of false negatives. In a full WES/WGS analysis pipeline, the false positive rate can be controlled by other strategies available for data analysis such as phenotype-based prioritization.^50–52^ For instance, combining the predictions of SQUIRLS with linkage analysis, candidate gene lists, or phenotype analysis would be likely to further improve rankings of causal variants.^50,51^

Many resources for genomic diagnostics precalculate scores for some subset of all possible variants. For instance, dbNSFP collects functional predictions and annotations for over 80,000,000 human nonsynonymous single-nucleotide variants and splice-site variants from various other algorithms that precompute values for all possible nucleotide changes in specified regions.^53^ However, this approach does not scale well for the prediction of splicing-relevant variation, which can affect multiple nucleotides and be located at arbitrary intronic and exonic positions. In our study, three of the approaches we compared with SQUIRLS offer precomputed scores but did not cover all tested variants. Of the 243 test variants, CADD missed 3 (1%), SpliceAI missed 27 (11%), and S-CAP missed 108 (43%). For clinical use, it is therefore important to optimize not only recall and precision but to engineer software such that it can analyze a wide range of variants in little time.

A limitation of SQUIRLS and all other approaches for computational prediction of SAVs in WES/WGS data that we are aware of, is that the algorithms predict the existence of an alteration of splicing, but do not attempt to predict the exact defect. In general, SAVs can be associated with a range of splice defects such as exon skipping, partial loss of exonic sequence, complete or partial intron inclusion, the creation of pseudoexons. Other investigations such as RNA-seq or target reverse-transcriptase PCR experiments are necessary to characterize these effects.

The UK 100,000 Genomes project and many other initiatives are poised to make genomic medicine part of healthcare for individuals with rare and common disease. In order to maximize the diagnostic yield of these programs, speed, efficiency, and ease of use are critical for technical incorporation of an algorithm into the diagnostic pipeline. However it is also crucial that the output of the algorithm is easily interpretable by the clinical scientists receiving the results of this pipeline in order that they can apply their findings to the treatment of the patient. In this work, we have presented an accurate and interpretable algorithmic approach for analyzing non-canonical splice variants that to date have been difficult to assess in exome or genome data. SQUIRLS combines state of the art accuracy with the ability to analyze arbitrary variants. On typical mid-range consumer hardware, SQUIRLS can analyze an exome file within a minute. To our knowledge, SQUIRLS is currently the only software that combines these abilities.

## Funding

This work was supported by the Horizon 2020 research and innovation program SOLVE-RD (grant 779257). Additional funding was provided by Monarch R24 [2R24OD011883-05A1] and by the National Scholarship Program of the Slovak Republic.

## Web resources

SQUIRLS download, https://github.com/TheJacksonLaboratory/Squirls/releases

SQUIRLS source code, https://github.com/TheJacksonLaboratory/Squirls

SQUIRLS manual, https://squirls.readthedocs.io/en/latest/

## Notes

### Competing Interest Statement

The authors have declared no competing interest.

