## Supplementary material for "Interpretable prioritization of splice variants in diagnostic next-generation sequencing": Suppl. Figures

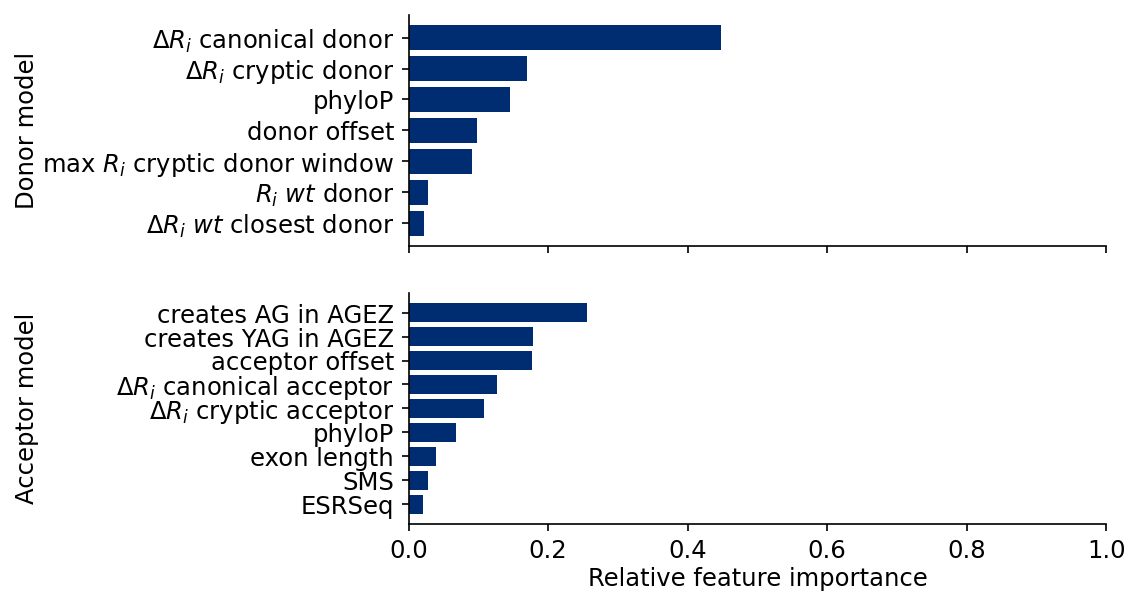

**Figure S1** - Relative feature importances in the donor/acceptor random forest classifiers. If the feature is used at the top of a decision tree, then it contributes to the prediction of a larger proportion of variants.

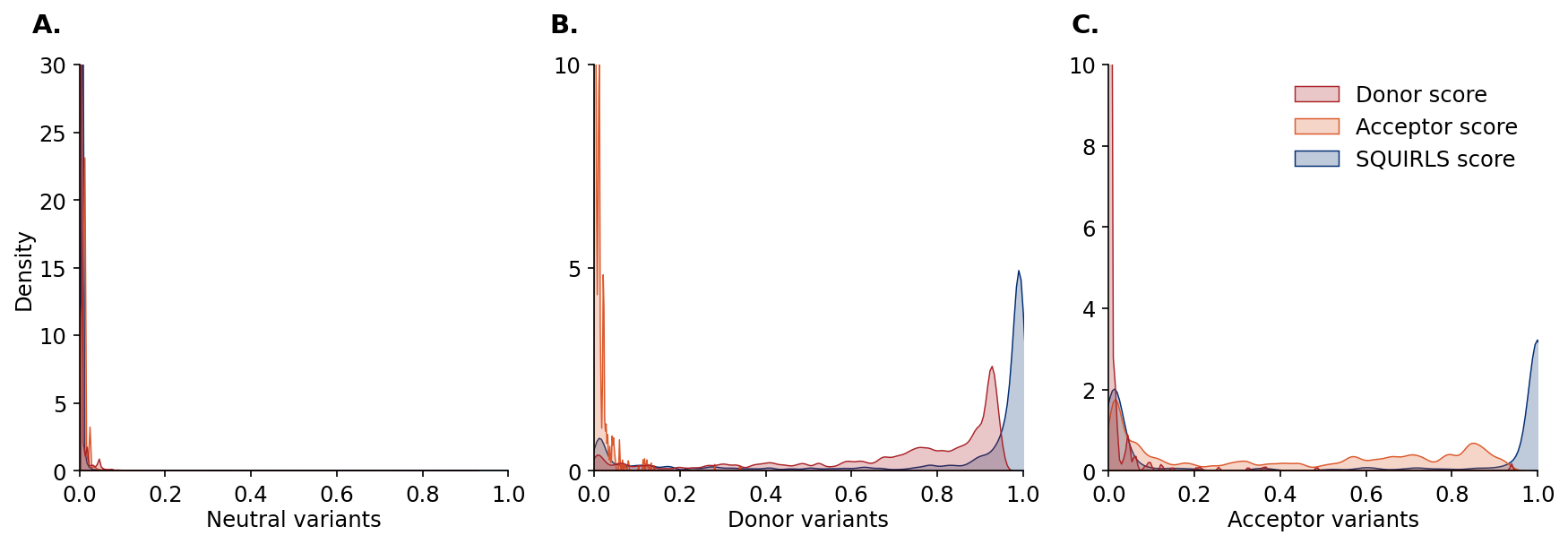

**Figure S2** - Density estimate for donor and acceptor scores calculated for all non-canonical training set variants (n_donor_ = 1,139, n_acceptor_ = 484, n_neutral_ = 62,986). **A)** SQUIRLS assigns low donor, acceptor, as well as the final SQUIRLS score to 62,986 splicing neutral variants. **B)** Unscaled scores for donor variants (generated by the donor-specific random forest classifier). **C)** Unscaled acceptor scores for acceptor variants (generated by the acceptor-specific random forest classifier). The models show site-specificity (e.g., donor variants are not assigned high acceptor scores and acceptor variants are not assigned high donor scores). The raw scores from the donor and acceptor random forest classifiers do not span the entire range of **[**0,1**].** SQUIRLS uses logistic regression to generate the final SQUIRLS score.

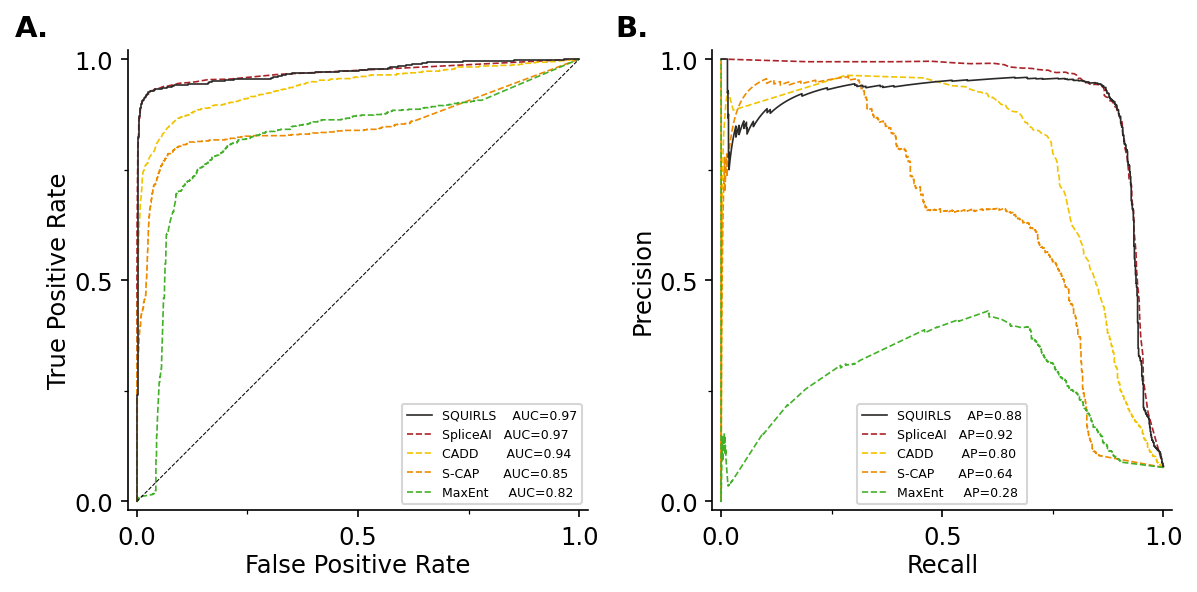

**Figure S3** - Performance of SQUIRLS, SpliceAI, S-CAP, CADD, and MaxEnt on all variants in the test set, including 6,008 canonical SAVs. A) Receiver operating characteristic curves indicate that SQUIRLS and SpliceAI achieve comparable performance. B) Precision-recall curves show that SQUIRLS and SpliceAI are able to find the most of the true splice variants, while maintaining high precision.

| 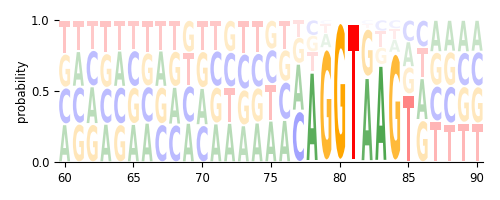 | 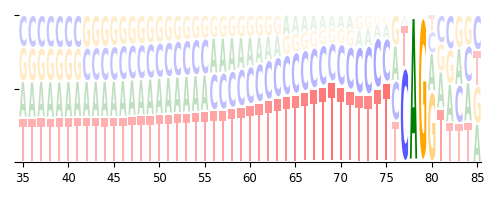 |
| --- | --- |
| 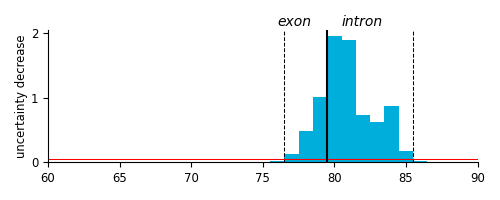 | 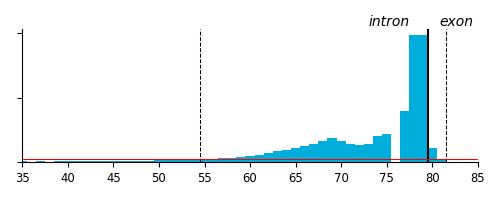 |
| **Figure S4** - Defining computational models of donor and acceptor sites. We aligned sequences neighboring splice junctions and determined probabilities of observing a base b at l-th position of the A) splice donor and acceptor B) sites. Probabilities are depicted as sequence logos where height of a character representing a base b corresponds to probability of observing b at l-th position of the splice junction.  After computing **R**_iw_ (methods), we summed the elements by columns to get the uncertainty decrease at l-th position of the splice donor C) and acceptor D) sites. We chose a heuristic threshold t=0.05 bits (red line) to correct for the background noise and we determined the size of splice donor and acceptor sites to be 9bp and 27bp, respectively. The splice site regions are denoted by the dashed vertical lines. | |

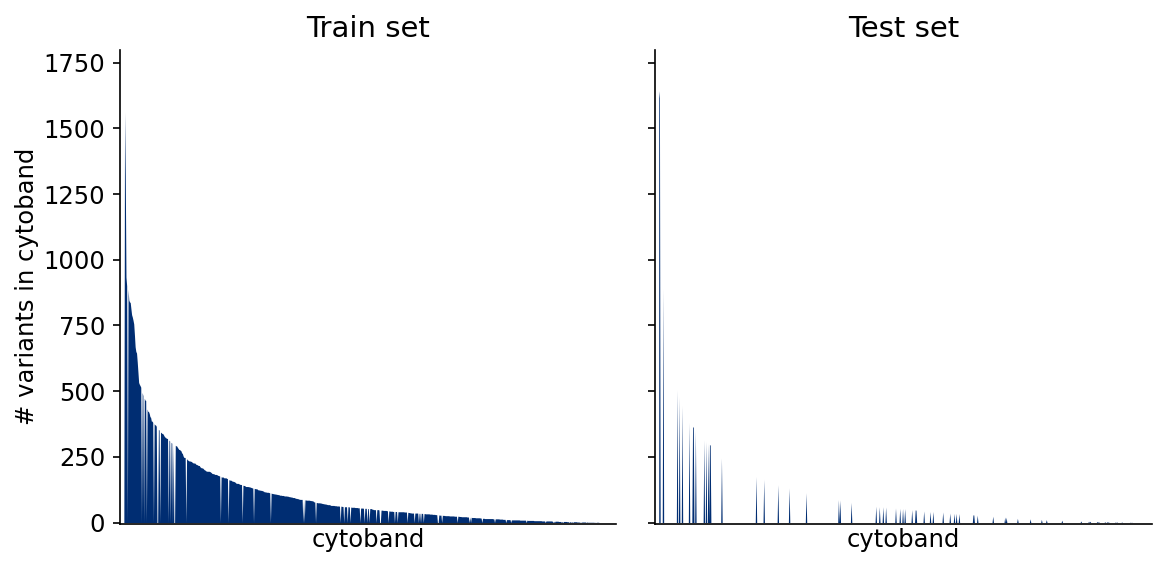

**Figure S5** - Cytoband-aware splitting of variants into training and test set. Each vertical line represents a cytoband, the line height represents the number of variants present within the cytoband.

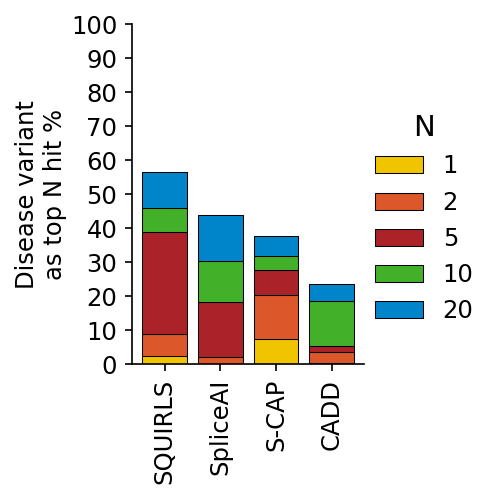

**Figure S6** - **Rank analysis simulation results.** 243 cases were analyzed with SQUIRLS, whereby a disease-associated SAV was spiked into a VCF file. The figure displays the same results as panels B and C of Fig. 5 of the main manuscript.
